# TipQuant: A robust algorithm for quantitative analysis of spatiotemporally dynamic activities in tip-growing cells

**DOI:** 10.64898/2026.05.20.725474

**Authors:** Jingzhe Guo, Julian Le Gouic, Rémi Rosenthal, Ailing Zou, Xiang Zhou, Nicolas Brunel, Zhenbiao Yang, Xinping Cui

**Affiliations:** Institute for Integrative Genome Biology and Department of Botany and Plant Sciences, University of California, Riverside, California 92521, USA; Capgemini Invent, 92130 Issy-les-Moulineaux, France; Quantmetry, l 52, rue d’Anjou, 75008 Paris, France; Metabolomics Research Center, Haixia Institute of Science and Technology, Fujian Agriculture and Forestry University, Fuzhou 350002, China; State Key Laboratory of Quantitative Synthetic Biology, Shenzhen Institute of Synthetic Biology, Shenzhen Institutes of Advanced Technology, Chinese Academy of Sciences, Shenzhen 518055, China; ENSIIE & Laboratoire de Mathématiques et Modélisation d’Evry, Université Paris Saclay, 91025 Evry, France; Institute of Emerging Agricultural Technology, Shenzhen University of Advanced Technology, Shenzhen, Guangdong, China; Faculty of Synthetic Biology, Shenzhen University of Advanced Technology, Shenzhen, Guangdong, China; Institute for Integrative Genome Biology and Department of Statistics, University of California,Riverside, California 92521, USA

## Abstract

Cell polarity, essential for cell development and function, relies on dynamic subcellular distribution of structural and signaling molecules. Tip growth, an extreme form of polar growth, involves unidirectional expansion at the apical region of cells and requires precise spatiotemporal coordination to achieve periodic and directional growth. Understanding their spatiotemporal dynamics is critical for elucidating mechanisms and functions of cell polarity. However, manual quantification of such dynamics is extremely time-consuming, hindering advancements in the field. Current algorithms have limited power and flexibility in analyzing the distribution and dynamics of molecules and structures, particularly for tip-growing cells with oscillatory and dynamic behavior. To address this challenge, we present TipQuant, an automated analysis tool that robustly detects tips and analyzes spatiotemporal dynamics of fluorescently labeled molecules/structures on plasma membranes and in cytoplasm at apices of tip-growing cells, enabling quantitative understanding of signaling and structural components in these systems.

## Introduction

Cell polarity, a fundamental asymmetric property of the cell, is essential for its development and function, such as cell differentiation, movement, morphogenesis, and polar and directional growth. Cell polarity can be found in all cellular organisms. In all cases, the formation of cell polarity relies on the dynamic distribution and changes in the type and quantity of subcellular structures and molecules that are typically interconnected into a network at the polar site (Drubin & Nelson, 1996; Yang, 2008; Chiou *et al*., 2021). Understanding their spatial distribution over time (i.e., spatiotemporal dynamic) is critical for elucidating the mechanism and function for the formation of a specific cell polarity. Manual quantification of these dynamics is extremely time-consuming, hindering progress in the field and limiting the integration of mathematical modeling into studies of cell polarity formation. Thus, computer-aided robust automated analyses are needed to advance the field, but are challenging, especially in a cell system with moving or changing sites of polarity, which is quite common in moving, growing, and dividing cells.

Tip growth is an extreme form of polar growth characterized by a unidirectional expansion in the apical region of the cell and is found in many cell types throughout the eukaryotic kingdom, such as pollen tubes and root hairs in plants, hyphae/mycelia in fungi, and neuronal axons in animals (Yang, 2008; Qin & Yang, 2011). Pollen tube tip growth is required for the delivery of sperms within the female tissues and is needed for plant reproduction. Hyphal growth is essential for the colonization and invasion of fungi in the host tissues, while axon extension is necessary for neuronal development and function. Tip growth generates a highly elongated cell with a cylindrical shape and involves the expansion of the cell only in the dynamic apical region of the cell where membrane and extracellular matrix materials are inserted through exocytosis. The process in different systems is commonly controlled by a highly dynamic and complicated network of signaling molecules and structures centered on the Rho-family GTPases. Hence tip-growing cells provide an excellent system for studying spatiotemporal coordination of dynamic cellular activities.

In particular, the pollen tube serves as an excellent model system for investigating such spatiotemporal coordination and the mechanisms underlying tip growth, because it can be conveniently manipulated and imaged *in vitro* and various genetic and molecular tools are available for this system. Pollen tube tip growth may oscillate rapidly with periods as short as 5-6 seconds (Guo *et al*., 2022). *In vivo* pollen tube growth is directional, guided by attractive signals secreted from the female tissues (Higashiyama *et al*., 2001; Okuda *et al*., 2009; Luo *et al*., 2017). As in other walled tip-growing cell types, the growth of pollen tubes requires the expansion of the cell wall at the apex driven by internal turgor pressure, while the mechanical properties of the cell wall determine the morphology of the cell (Hepler *et al*., 2013). To achieve periodic and directional growth, the deposition of the cell surface materials is controlled precisely in space and time by a spatiotemporally coordinated signaling network that is centered on ROP (**R**h**o**-like GTPase from **p**lants) signaling and composed of multiple downstream pathways, feedforward and feedback loops, cytoskeletal dynamic, and influxes and effluxes of ions such as calcium (Yang, 2008; Hwang *et al*., 2010; Qin & Yang, 2011; Luo *et al*., 2017; Tian *et al*., 2019).

This network is characterized by its remarkable spatiotemporal dynamics (Cheung & Wu, 2008; Zonia, 2010; Guan *et al*., 2013). Spatially, it is highly polarized: dense populations of exocytic vesicles, supported by longitudinal actin filaments and an apical actin meshwork, move towards the extreme apex of pollen tubes, delivering cell membrane and cell wall components to the tip, while endocytosis retrieves excessive materials from the subapical or apical regions (Guo, J. & Yang, Z., 2020). Active ROP1 and its associated signaling components, localized to the apical membrane, play an essential role in maintaining cell polarity by regulating actin dynamics and exocytosis (Fu *et al*., 2001; Gu *et al*., 2005; Lee *et al*., 2008; Guo, J. & Yang, Z., 2020). Calcium ions also show a tip-high gradient and are crucial for pollen tube growth by participating in multiple pathways including the ROP signaling (Pierson *et al*., 1996; Li *et al*., 1999; Yan *et al*., 2009; Guo *et al*., 2022). Temporally, this system is oscillatory and extremely dynamic: ROP1 activity, calcium concentration, and accumulation of vesicles at the cell tip exhibit rapid and correlated oscillations resulting in oscillating growth (Hwang *et al*., 2005; Lee *et al*., 2008; Yan *et al*., 2009). The mechanisms underlying the polar growth of other tip-growing cells, such as pollen tubes, root hairs, and fungal hyphae, are evolutionarily conserved, and this growth can be mediated by different protein families or even different organizing structures, such as the Spitzenkörper, depending on the cell type and organism (Qin & Yang, 2011). Therefore, the study of polar growth in these cell models relies heavily upon the analysis of the distribution and dynamics of various molecules and structures, yielding large amounts of video image data that require quantification and interpretation, which is extremely time-consuming by manual measurements.

Available algorithms or codes for analyzing tip-growing cells have limited power and flexibility for analyzing the spatiotemporal dynamics of labeled molecular or structural events. For example, the CHUKNORRIS, an algorithm for detecting oscillations of ion fluxes as well as growth rate data, rely on a user generated kymograph, therefore, has the element of user-subjectivity and contains partial spatiotemporal information of molecular components along the central axis of the cell (Damineli *et al*., 2017). The Automated Stack Iterative Subtraction (ASIST) method (Ponvert *et al*., 2019) and the contour-based coordinate normalization (CCN) method (Kang *et al*., 2023) have been successfully applied to analyze growth parameters of tip-growing cells, such as pollen tubes, root hair, and zygotes. However, their analyses only limited to growth rate and directionality while unable to analyze spatiotemporal distribution of molecules and structural components. The TIGRMUM toolkit, detecting the tip of pollen tubes by weighting the ellipse method and morphological thinning (Burri *et al*., 2020), perform measurement of fluorescence intensity or ratio of two fluorescent channels over a fixed cytosolic region of interest (ROI) with capability of analyzing the left and right half ROIs separately. AMEBaS (Automatic Midline Extraction and Background Subtraction of Ratiometric Fluorescence Time-Lapses of Polarized Single Cells) generate kymograph along the midline of tip-growing cells lacking quantitative measurement over any user selectable ROIs (Badain *et al*., 2023). A more recently developed KymoTip allows users to analyze the growth behavior of plant tip-growing cells and generate centerline kymographs for various fluorescent markers; however, it lacks functionality for quantifying intensity changes over time (Kang et al., 2026). We attempt to address this problem by initially developing a TipQAD algorithm to automatically quantify the spatiotemporal dynamics of fluorescently labeled molecules or structures on the plasma membrane (PM) and in the cytoplasm at the apex of tip-growing cells (Tambo *et al*., 2020). In the TipQAD algorithm, the contour of a tip-growing cell is first fitted with ellipses, the intersection of the PM and the major axis of the best-fit ellipse is determined as the growing tip. While the TipQAD has provided better analyzing power over the previously developed methods, it suffers from several key issues: First, the tip determination highly depends on the shape of the growing cells therefore unable to analyze cells that change shapes dramatically such as large changes in growth direction or cell diameter. Second, it only analyzing a fixed horse-shoe shaped ROI in the cytoplasm, lacking the flexibility of ROI choices for quantitative analyses.

Here, we introduce TipQuant (<u>Tip Quant</u>ification), a Python-based automated tool designed for detecting tips and analyzing the spatiotemporal dynamics of fluorescently labeled molecules and structures on the plasma membrane (PM) and in the cytoplasm at the apex of tip-growing cell. We have demonstrated the efficacy of TipQuant in correlating the spatiotemporal dynamics of two molecular events - ROP and PM Ca^2+^ influxes - within the same cell. Consequently, TipQuant proves to be an essential resource for quantitative analysis of spatiotemporal dynamics of signaling and structural components in tip-growing cells. It also serves as a valuable data source for mathematical and computational modeling studies aimed at a quantitative understanding of these dynamics.

## Results

### Algorithm development

Central to the automated analyses of tip-growing cells is robust detection of growing tips, especially when pollen tubes have complicated growth patterns such as turning and zigzagging. Tip growth is defined by directional deposition of membranes and/or cell wall materials, such that a directional expansion of the cell is accompanied by an observable morphological change. Therefore, we developed a tip detecting algorithm which defines the growing tip as the site with maximum expansion. Specifically, for a given video serial, the algorithm first performs contour segmentation of a tip-growing cell on all frames (see **Supplemental Method S1** for details of contour segmentation), then detects the growing region by identifying maximum spatial dislocation of cell contour between two frames parted with a user-defined step (FRAME _n_ versus FRAME _n + t_) (Figure 1A). During the tip detection, we create a sliding window (in user-defined pixel numbers) moving along the expanded contour pixel by pixel, we then calculate the curvature of the window and create a normal vector for each sliding window. Finally, we calculate the displacement of each sliding window along its normal vector and define the center of the sliding window with the maximum displacement as the tip (Figure 1B, also see **Supplemental Method S2** for details of tip detection). Based on the algorithm-recognized contour, including spatial coordinates of contour and tip position, we further developed codes for measuring mean fluorescence intensity on user-defined PM and cytosolic region of interest, as well as PM kymograph (see detailed coding information in **Supplemental Method S3**).

**Figure 1.**
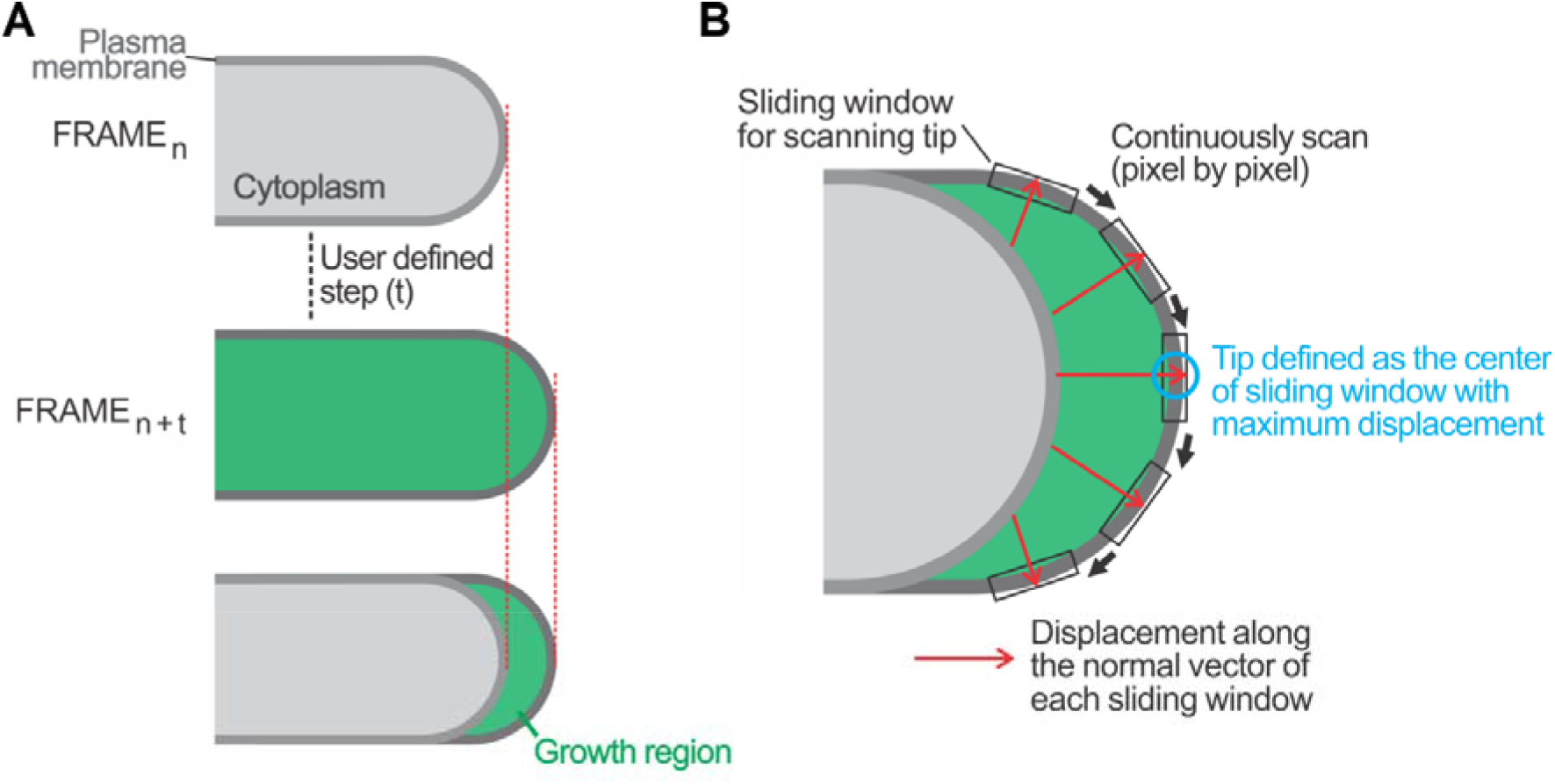
Simplified schematic depicting the process of tip detection. The tip detection involves two steps: growth region detection (A) and tip determination on the plasma membrane (B). In the first step (A), the growth region is identified by the spatial dislocation of cell contour between two frames separated by a user-defined step (FRAME *_n_* versus FRAME *_n + f_*). In the next step (B), a sliding window (of user-defined pixel numbers, depicted by the black rectangle along the PM) moves along the contour pixel by pixel; a normal vector is then computed for each window after calculating its curvature. Subsequently, the displacement of each sliding window along its normal vector (illustrated by the red arrow) is determined, and the window with the maximum displacement is defined as the tip.

### TipQuant robustly segments and detects growing tips

To validate the performance of our tip detection algorithm, we challenged the code with two types of pollen tubes with complicated growth patterns: pollen tubes forced to become narrower when squeezing through a narrow microfluid channel (Zhou *et al*., 2021) and pollen tubes turning to a female attraction signal AtLURE1 (Luo *et al*., 2017). In the first case, a pollen tube expressing the ROP activity sensor CRIB-GFP is growing from a wide-channel (9 µm in width) to a narrow channel (4 µm in width) inside a TipChip microfluid device (Zhou *et al*., 2021) (Figure 2A). With a typical diameter of 6 µm, the diameter of the pollen tube was forced to 4 µm to squeeze through the narrow channel (Figure 2B). Nevertheless, our code still robustly segregates and detected the growing tip throughout the shape changing process (Figure 2C). In the second case, we ask the code to track the growing tip of a pollen tube turning toward the AtLURE1 attraction peptides (Figure 2D). Although the pollen tube has made several turns and grown into different focal planes compared to the first few frames, our code still manages to track the growing tip (Figure 2D). We also have successfully segregated and tracked the growing tip of fungal hyphae cells (see later results), indicating the wide applicability of our code to various tip-growing cells. These results demonstrate the robustness of our code for segmentation and tip detection for tip-growing cells regardless of their growing behavior.

**Figure 2.**
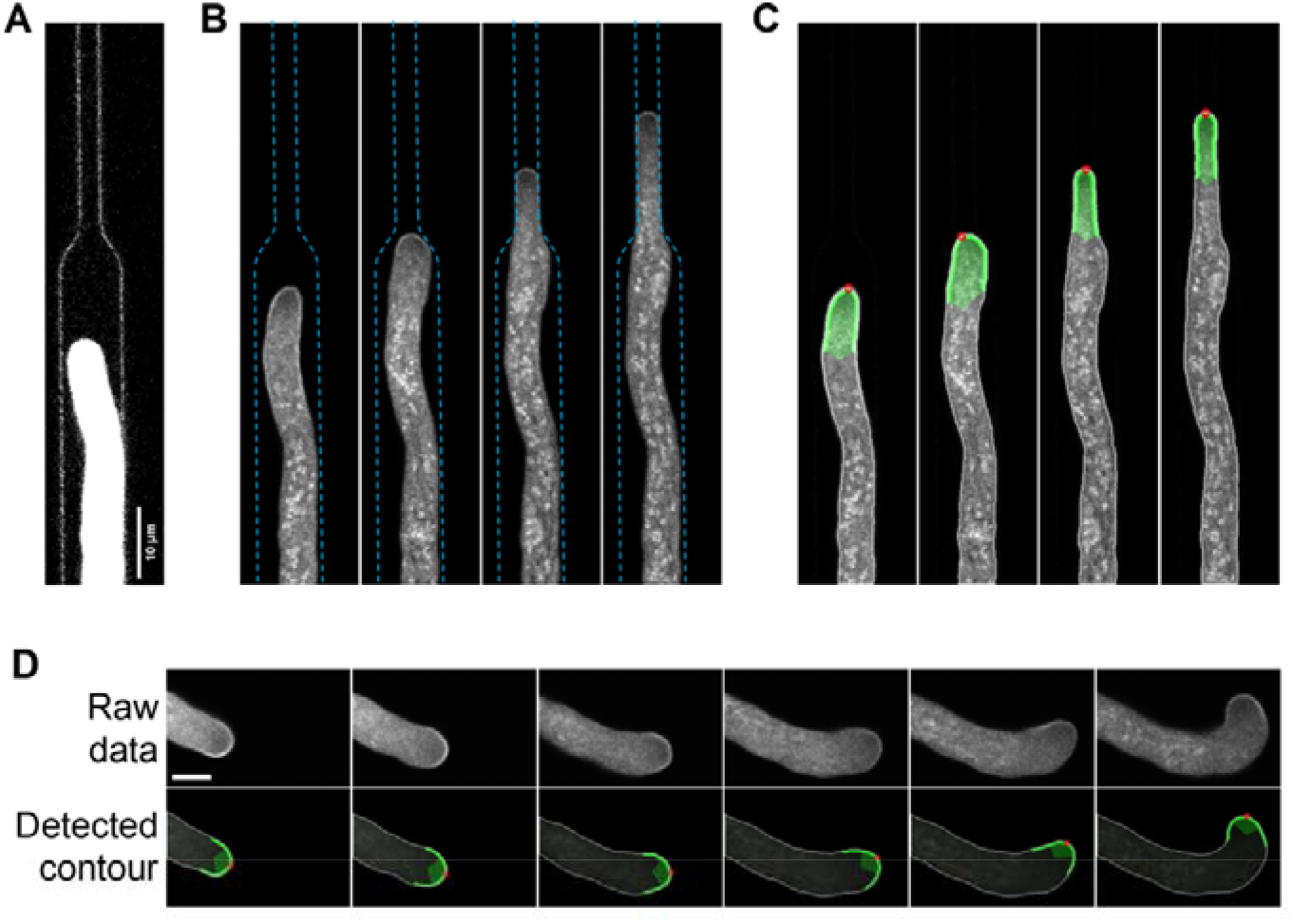
Dynamic segmentation and tip detection of tip-growing cells with substantial morphological changes. **A-C**. Tip detection of a CRIB4-GFP pollen tube growing on a TipChip microfluidic channel. The channel comprises a wide part and a narrow part (shown in **A** by enhanced image contrast), and the pollen tube growing in the wide channel is forced to squeeze into the narrow channel (raw image shown in **B** with the contour of the channel highlighted by the blue dotted line). The output of the TipQuant-detected pollen tube is shown in **C**. Scale bar is 10 μm. **D**. Tip detection of a CRIB4-GFP pollen tube showing frequent turning behaviors attracted by manually applied pollen tube attractant AtLUREI. The upper panel shows raw images, and the lower panel shows the TipQuant-detected pollen tube. Scale bar is 5 μm. In all TipQuant-detected images, the tip is marked with red circle, the PM and cytosolic regions at the tip are highlighted with brighter green lines and lighter green shades respectively.

### Quantitative measurement on user-defined ROIs and PM kymograph generation

Quantifying spatiotemporal changes in molecule distribution is crucial for understanding the mechanisms regulating tip growth. As previously mentioned, current methods for analyzing tip-growing cells either lack the capability or have limited power to generate kymograph and perform quantitative measurements over region of interest (ROIs). The TIGRMUM toolkit is the only code that generates kymograph along the central line of pollen tubes and measures fixed tip ROIs (Burri *et al*., 2020).

In contrast, our TipQuant code offers flexible options for selecting ROIs, with three different user-adjustable shapes of ROIs within the cytoplasm and on the PM for quantitatively analyzing mean fluorescence intensity. Additionally, TipQuant can analyze fluorescence intensity distribution on user-defined PM regions and generate a 2D kymograph of the spatiotemporal distribution of labeled molecular or structural components on the PM.

#### Quantitative measurement on user-defined cytosolic ROIs

Quantifying cellular and signaling components, which often have distinct subcellular localizations, requires highly adaptable ROIs to accommodate their unique spatial distribution patterns. To address this need, our TipQuant code offers three types of cytosolic ROIs for measuring signal intensities: a V-cone shaped region (ROI-A), a whole cytosolic region from tip to shank (ROI-B), and a circular region within the cytoplasm (ROI-C) (Figure 3A). These ROIs are adjustable in both position and size, making them suitable for analyzing a wide range of proteins with diverse subcellular localizations. Specifically, the V-cone-shaped ROI-A is ideal for proteins involved in polar secretion, the whole cytosolic ROI-B is suitable for most cytosolic proteins and reporters of various ions or signaling molecules, and the circular ROI-C within the cytosolic tip is designed for specific subregions, such as the characteristic Spitzenkörper in fungal hyphae.

**Figure 3.**
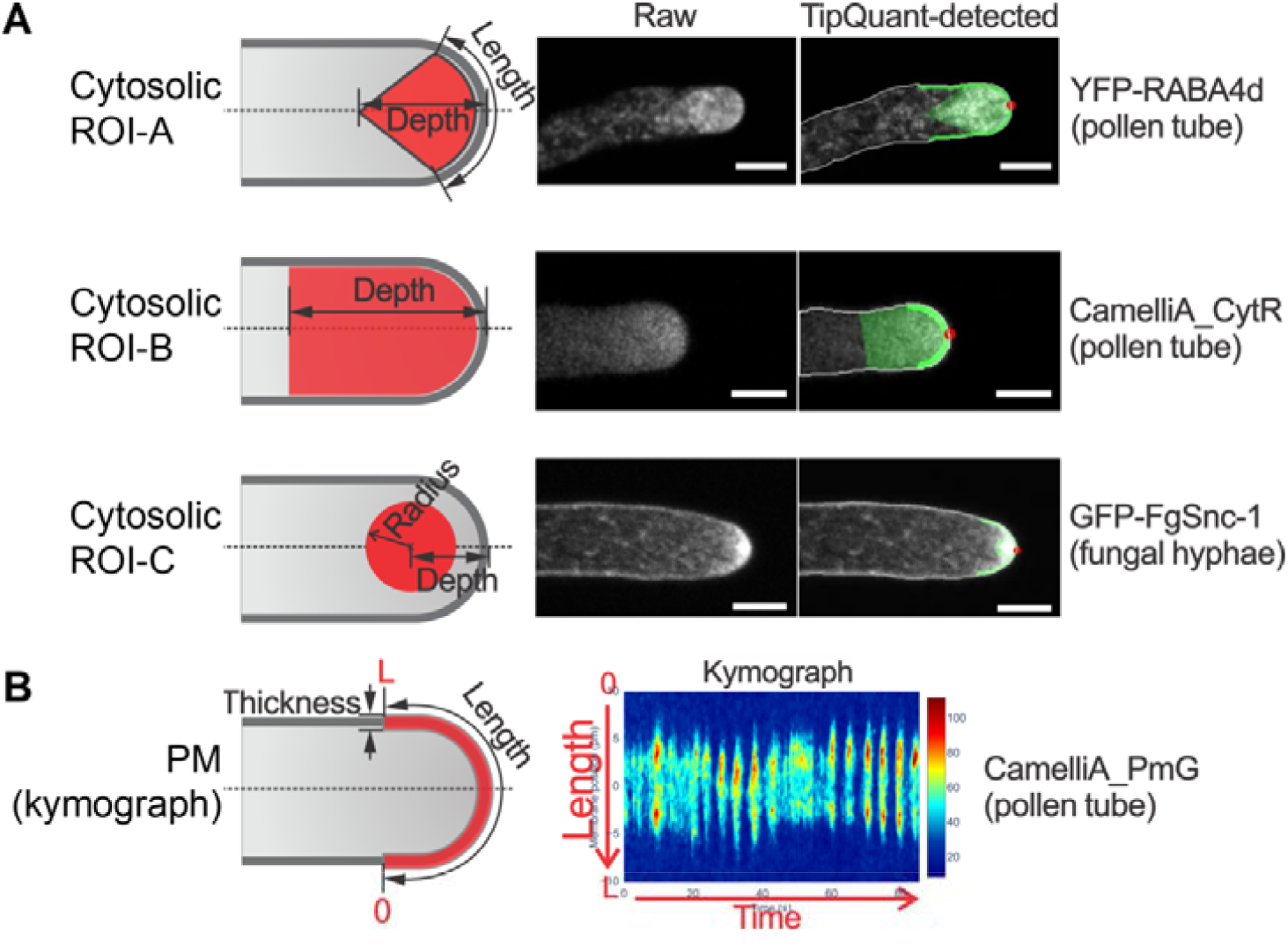
TipQuant enables quantitative analyses over versatile selections of ROIs. **A**. The code allows measuring the mean fluorescence intensity of cytosolic ROIs of a V-cone shaped region (ROI-A), whole cytosolic region from tip to a user-defined depth (ROI-B), and circular region within the cytoplasm (ROI-C). All three types of cytosolic ROIs have user adjustable parameters to fit the subcellular pattern of interest proteins or dyes (left column). The middle and right columns show the representative raw images of the tip-growing cells (middle column) and the TipQuant-detected results (right column) with light green shades highlighting the cytosolic region for measurement. Scale bars are 5 μm. **B**. Mean fluorescence intensity of a PM region spanning evenly from the tip with a user-defined length and thickness (left panel) along with a kymograph (right panel) generated by the TipQuant.

#### Quantitative measurement on user-defined PM ROI and PM kymograph generation

The plasma membrane (PM), as the primary interface between the cell and its environment, hosts numerous signaling proteins, including receptors, ion transporters, and GTPases etc., which orchestrate cellular activity in response to internal and environmental cues. These signaling events often trigger changes in ion influxes, enrichment/depletion of PM-localized proteins, or the formation of protein subdomains on the PM. To better understand these cellular processes, the TipQuant code includes functions for spatial and temporal quantification on the PM by providing measurements of mean fluorescence intensity and generating kymograph. By accurately tracking the extreme tip of the PM, the TipQuant code allows users to measure fluorescence intensity distribution along a user-defined PM region centered around the apex, which is then converted into kymograph slices for each time frame. A complete PM kymograph is generated by aggregating these slices, providing a dynamic view of the PM activity over time, along with the mean PM fluorescence intensity.

### TipQuant-generated quantitative data faithfully represent the ground truth

The correctness and coherence of the TipQuant methodology are evaluated by comparing the TipQuant output with manually measured data using ImageJ, which serves as the ground truth (GT). The mean fluorescence intensity measurements over time across three distinct cytosolic ROIs are compared with their respective GT data (Figure 4A-D). All comparisons demonstrate a high level of consistency, as indicated by the strong Pearson’s correlation (Figure 4E).

**Figure 4.**
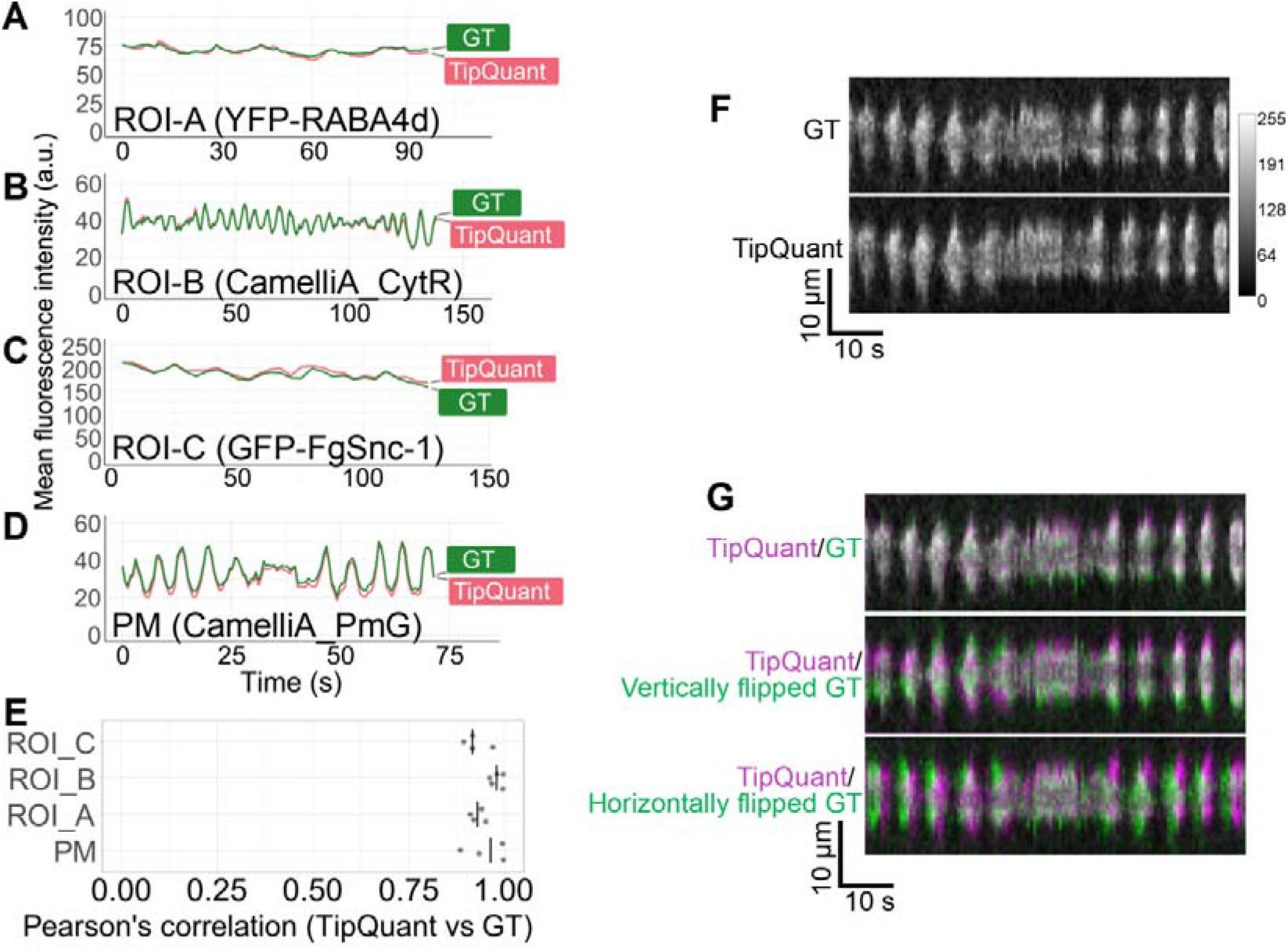
Validation of TipQuant-generated quantitative data and kymographs. **A-D**. Representative plots of ground truth (GT) and TipQuant-generated mean fluorescence intensity changes with time in cytosolic ROI-A (A), ROI-B (B), ROIC (**C**), and PM (**D**). The corresponding marker lines are indicated in parentheses following each ROI label. **E**. Plot of Pearson’s correlation of the GT and TipQuant-generated data in the above measured ROIs. Each dot represents comparison within an independent sample. **F-H**. Validation of the PM kymograph of CamelliA_PmG generated by manual measurement (GT, ground truth) and TipQuant (**F**). Composite images of purple kymograph generated by TipQuant superimposed with green-colored GT kymograph **(G**, top panel) or vertically flipped and horizontally flipped GT kymograph (G, middle and lower panels respectively). The white color indicates perfect match between the two compared kymographs, while the more distinct non-overlapping purple and green colors, the poorer the match.

We also compared the TipQuant-generated PM kymograph with manually curated data. The manually generated kymograph was created by vertically stacking intensity distributions measured over a 20 µm PM region flanking the tip apex at each time frame. Due to human subjectivity, the determination of the tip apex was arbitrary and not consistent with the apex determined by TipQuant. Therefore, a direct comparison between the two kymographs is not feasible. To address this, we first realigned the manually generated kymograph with the TipQuant-generated kymograph (Figure 4F, lower panel) by shifting the Y-coordinate of each PM slice to align with the TipQuant data (Figure 4F, see Realignment of ground truth PM kymograph with TipQuant-generated kymograph in the Material and Methods section.). We then compared the two kymographs by superimposing the color-encoded kymographs (purple for TipQuant-generated and green for the manually curated kymograph, therefore two matched patterns generate white color when superimposed.) using ImageJ software (Figure 4G). To further validate the accuracy of the TipQuant-generated kymograph, we performed the same superimposition process between a vertically or horizontally flipped manually curated kymograph and the TipQuant-generated kymograph. The results indicate a high degree of correspondence only between the TipQuant-generated kymograph and the manually curated data. (Figure 4G).

### TipQuant revealed temporal relationship between oscillations of PM ROP activity and Ca^2+^ influxes

After validating the fidelity of the TipQuant-generated measurements, we examined the potential of TipQuant to improve our understanding of signaling events in cellular processes. Using the ROP GTPase activity reporter CRIB4-GFP(Luo *et al*., 2017) and the PM Ca^2+^ influx reporter CamelliA_PmR(Guo *et al*., 2022) in Arabidopsis pollen tubes, we quantified temporal changes in these two key signaling components at the PM with TipQuant. Figure 5 presents a representative pollen tube expressing both reporters, together with TipQuant-based quantification of the PM ROP activity and Ca^2+^ influx using the mean fluorescence intensity at the PM as readout. Spearman’s correlation analysis of video data from 12 independent pollen tubes revealed a strong positive correlation between oscillations in the ROP activity and PM Ca^2+^ influx (Figure 5C, Supplemental Figure S1)

**Figure 5.**
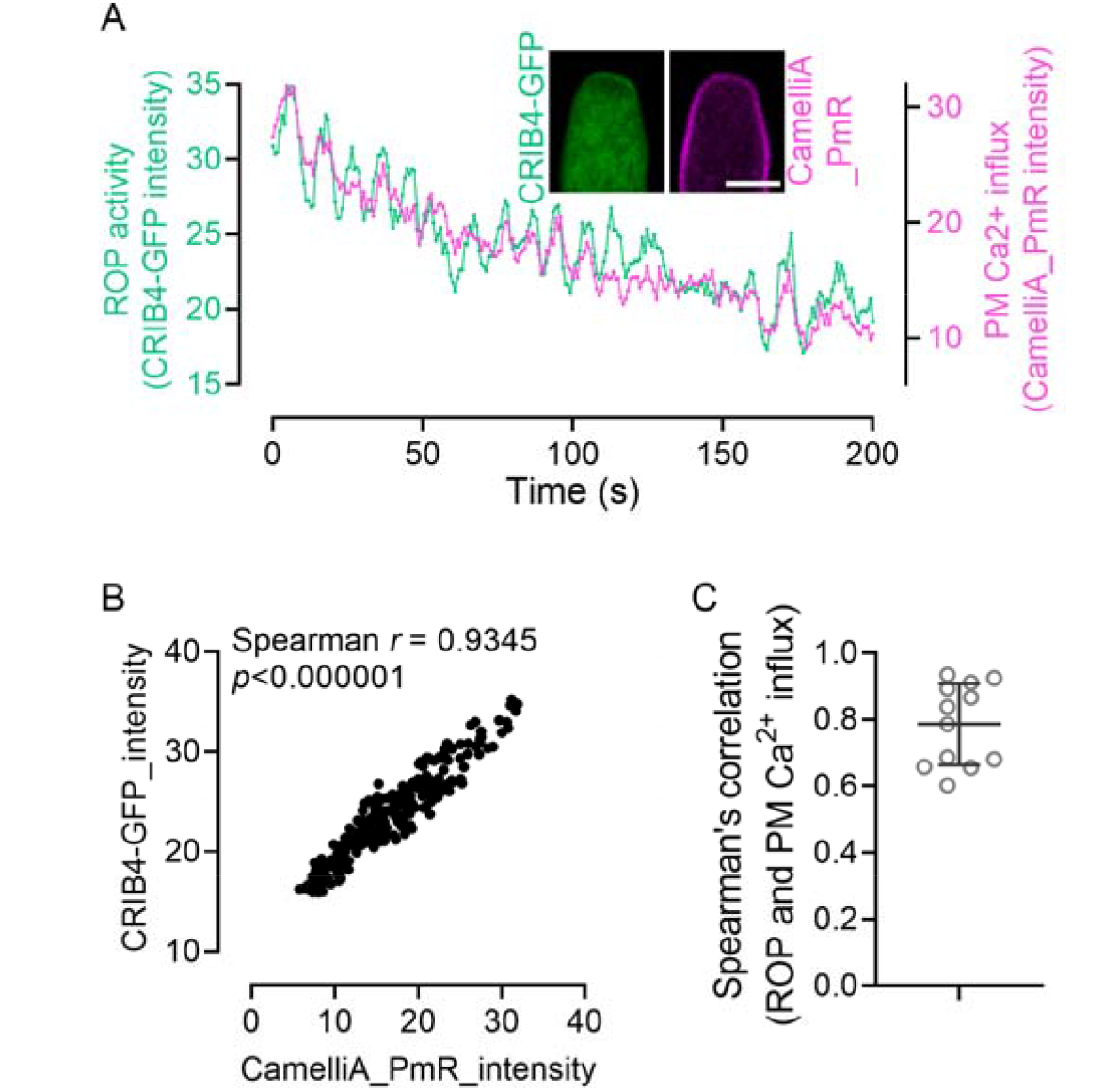
Simultaneously analyses of ROP activity and PM Ca^2+^ influx in pollen tubes. **A**. Time-course plot showing mean PM fluorescence intensity changes for the ROP activity sensor CRIB4-GFP and the PM Ca^2+^ influx sensor CamelliA_PmR. Insets show representative images of the pollen tube expressing CRIB4-GFP (left) and CamelliA_PmR (right). **B**. Scatter plot of paired CRIB4-GFP and CamelliA_PmR intensity values at each time point, with Spearman’s correlation coefficient and *p*-value indicated at the top. **C**. Scatter plot of the Spearman correlation coefficients (*r*) between CRIB4-GFP and CamelliA_PmR intensities from videos of 12 different pollen tubes. Scale bar is 5 μm in **A**.

## Discussion

We have developed and extensively tested the TipQuant code, demonstrating its ability to automatically and accurately analyze the spatiotemporal distribution of key molecular and structural components in tip-growing cells in an unbiased manner. It has shown exceptional capability in analyzing various user-defined ROIs, thereby facilitating the discovery of hidden information within existing confocal microscopy imaging datasets.

Our code detects growing tips by identifying the site with maximum spatial displacement, rather than relying on ellipse fitting as applied in several other reported codes(Burri *et al*., 2020; Tambo *et al*., 2020). Moreover, we offer users greater flexibility in selecting desired ROIs for confocal microscopy data analysis compared to other existing codes. Consequently, our code offers several distinct advantages over existing alternatives, as discussed below.

First, our TipQuant code can effectively process tip-growing cells with complex growth behaviors, as demonstrated by its ability to segment and track pollen tubes that undergo dramatic changes in diameter when growing into microfluid chamber and when navigating multiple turns toward the AtLURE peptide. However, we did notice that tip detection become less robust for cells that turn dramatically, such as turning more than 90° or making a U-turn. Second, our code offers maximum flexibility by allowing users to define ROIs on both the PM and in cytosolic regions for unbiased and automated data analysis. Lastly, our code generates advanced dataset by providing kymograph data along the PM, the primary interface for signaling transduction with the external environment, compared to traditional kymograph, which only display data along the central axis.

In addition, we tested our code on two different types of tip-growing cells - pollen tubes and fungal hyphae – using images acquired on major commercial confocal microscopy systems from Leica, Nikon, and Zeiss, with Nyquist sampling rates between 0.5x to 2x. These tests demonstrate the code’s promising applicability to various tip-growing cells and its compatibility with a wide range of confocal microscopy platforms.

Despite these advantages, the TipQuant algorithm is not applicable to non-growing cells, as its core detection mechanism relies on cell contour expansion between frames. However, quantitative measurements of non-growing cells can be readily performed using fixed ROIs in widely used image analysis software such as ImageJ (Schneider *et al*., 2012). Although uncommon, some tip-growing cells produce multiple tips(Jones *et al*., 2002; Wudick *et al*., 2018). Currently, TipQuant cannot process videos featuring multiple growing tips; however, users can overcome this by cropping the video to isolate a single growing tip prior to analysis. Additionally, TipQuant-generated quantitative data inherently lack information from the initial frames (determined by the user-defined step ‘*N*’ for cell expansion detection) due to the absence of reference frames for tip detection. Lastly, TipQuant currently works only with confocal imaging data, as it is not programmed to handle out-of-focus blur at cell contour boundaries in epifluorescence microscopy images.

## Material and Methods

### Computing platform

The TipQuant code requires Python 3.11 or later. While it should work with newer versions, Python 3.11 is recommended. All necessary dependent packages, as listed in the *environment*.*yml* file, will be automatically installed when the code is run. The code has been rigorously tested on macOS (a Macbook Pro M1 running macOS Ventura 13.4, and a MacBook Pro M5 running macOS Tahoe 26.3), and Windows (10, and 11) operating systems and has been shown to run smoothly.

### Video data

The video of Arabidopsis pollen tube expressing CRIB4-GFP squeezing through the narrow-gap region of a TipChip microfluid chamber was generated as described previously (Zhou *et al*., 2021).

The video of Arabidopsis pollen tube expressing CRIB4-GFP turning towards the attractant AtLURE1.2 were kindly shared by Dr. Nan Luo (Luo *et al*., 2017).

Videos of pollen tubes expressing CamelliA_PmG (PM Ca^2+^ influx) and CamelliA_CytR (cytosolic Ca^2+^ level), either separately or simultaneously, are derived from the authors’ previously published experimental observations (Guo *et al*., 2022) Videos of *Fusarium graminearum* hyphae expressing GFP-FgSnc1 were kindly provided by Dr. Wenhui Zheng (Zheng *et al*., 2021).

Videos of Arabidopsis pollen tubes expressing YFP-RABA4d were generated for this research. Briefly, pollen tubes expressing YFP-RABA4d driven by the pollen-specific promoter Lat52 (Szumlanski & Nielsen, 2009) were germinated on pollen germination medium and mounted for observation as previously described(Guo, Jingzhe & Yang, Zhenbiao, 2020). The fluorescence emission of YFP-RABA4d was collected between 525nm and 600nm while being excited by a 514nm laser under a Leica SP5II confocal microscope equipped with a Leica HCX PL APO CS 40x/1.10 WATER UV objective lens.

Videos of Arabidopsis pollen tubes co-expressing CRIB4-GFP and CamelliA_PmR were generated as follows. A homozygous Arabidopsis line expressing both markers was produced through genetic crossing. Pollen tubes of the dual-marker line were germinated, mounted as described above, and observed under Leica STELLARIS FALCON FLIM Microscope equipped with a Leica HC PL APO CS2 40x/1.10 water-immersion objective lens. The fluorescence emission of CRIB4-GFP was collected between 498nm and 570nm while being excited by a 488nm laser, and the emission of CamelliA_PmR was collected between 575nm and 700nm while being excited by a 561nm laser. Both fluorophores were imaged sequentially to avoid bleed-through.

### Generation of the ground truth data

We used Fiji/ImageJ to measure and generate the ground truth data for the mean fluorescence intensity of PM signals, various cytosolic ROIs and the PM kymograph.

#### Generation of the ground truth data of PM fluorescence intensity and kymograph

A 20 µm-long PM region at the apical tip of CamelliA_PmG pollen tube is manually traced using the *segmented line* tool. The fluorescence intensity of pixels in the selected region is then read out using the *plot profile* function, with the line width set to 5 pixels. This process is repeated for every frame of the video to collect the full dataset of PM fluorescence intensity distributions over time. This collection is further processed using Pandas (McKinney, 2010) and Scikit-fda Python packages (Ramos-Carreño *et al*., 2024) to generate the ground truth data for the PM kymograph. The mean PM fluorescence intensity over time is calculated as the average fluorescence intensity of the PM in each frame.

#### Generation of the ground truth data of fluorescence intensity in various cytosolic ROIs

Various cytosolic ROIs were manually traced using *Polygon selections* tool in each time frame, followed by measuring mean fluorescence intensity.

### Realignment of ground truth PM kymograph with TipQuant-generated kymograph

When comparing PM kymographs of CamelliA_PmG pollen tubes generated by manual measurement (ground truth, GT) and TipQuant (TQ), an important challenge in comparing the two intensities *I*^*GT*^*(x, t*) and *I*^*TQ*^*(x’,t)* is that they are not defined on the same spatial coordinate systems *x* and *x’*. Indeed, in both cases, the tip detection algorithms produce two different plane curves *C*^*GT*^ *= C*^*GT*^*(x)* and *C*^*TQ*^ (*x*) that can be parametrized by their respective arc lengths *x,x*^*’*^ (with total length *L*^*GT*^,*L*^*TQ*^). The coordinate *x* represents the distance along the curve *C*^*GT*^ between the point at location *x* and the left boundary *C*^*GT*^(0). The intensities *I^Gr^ I^TQ^* are functions defined on *C*^*GT*^ and *C*^*TQ*^ and we need to align them to ensure a fair comparison of the intensities. This requires finding a way to match every point *C*^*GT*^*(x)* to a unique point *C*^*TQ*^*(x’)* by determining the best function *h*. The function *h* establishes a one-to-one correspondence between the points of each tip: *h(C*^*GT*^*(x)) = C*^*TQ*^*(x’)* (and we simply write *h(x)=x’)*. This correspondence *h*: *C*^*GT*^ *-> C*^*TQ*^ is necessary to account for variations in the localization of the tip membrane. The simplest alignment corresponds to the case where the curves differ only by a translation by a factor *c*, and the two curves are matched with the function *h(x) = x + c*. The translation *c* is unknown and is selected to maximize the similarity between *I*^*GT*^*(x, t*) and *I*^*TQ*^*(x + c, t*) (assuming that *L*^*GT*^ *< L*^*TQ*^): the intensity functions must represent the same quantity.

In general, as the detected contours might involve a nonlinear transformation from one to the other, we need to consider more complex transformation *h*(*x*) for aligning *I*^*GT*^ (*x, t*) and *I*^*TQ*^ (*h*(*x*),*t*). The computation of an optimal alignment (warping) function *h*(*x*) can be achieved by minimizing the distance between the two functions *I*^*GT*^ (·,*t*) and *I*^*TQ*^(·,*t*). We consider the so-called Fisher-Rao distance for computing the best alignment, as it allows detecting similar shapes effectively. It is considered as one of the reference methods for alignment or registration of curves because it exhibits desirable theoretical properties like invariance to parametrization, and provides good results when used on real cases (Srivastava & Klassen, 2016). This distance involves the derivatives of the functions *I*^*GT*^ (·, *t*) and *I*^*TQ*^ (·,*t*) (since the derivative characterizes the peaks and valleys of *I*^*TQ*^ (·,*t*) more than *I*^*TQ*^ (·,*t*) itself), and we have used the implementation proposed in the Python Open Source Package scikit-fda (https://github.com/GAA-UAM/scikit-fda).

### Comparison ground truth and TipQuant-generated fluorescence intensity measurements

We use *corr* function in the MATLAB R2022a to calculate the Pearson’s linear correlation coefficient between the ground truth and TipQuant-generated fluorescence intensity vs. time data arrays.

## Acknowledgments

We would like to thank Dr. Nan Luo from the Yang lab for sharing the videos of Arabidopsis pollen tubes turning to AtLURE peptides, Dr. Wenhui Zheng from Fujian Agriculture and Forestry University for providing the videos of fugal hyphae. This research is partially supported by NSF DMS 1853698 awarded to Dr. Xinping Cui and Dr. Zhenbiao Yang, USDA Hatch Project AES-CE award (CA-R-STA-7132-H) to Dr. Xinping Cui.

## Author contributions

J.G. and Z.Y conceptualized the research; J. L. G., R. R., and N. B. developed the TipQuant codes with input from J. G.. J.G. conducted experimental observations, generated ground truth data, and wrote the original draft; A. Z. and X. Z. acquired videos of the pollen tubes co-expressing CRIB4-GFP and CamelliA_PmR, as well as pollen tubes passing through the microfluidic channel; N. B., Z. Y., and X. C. supervised the research and revised the manuscript; all authors have read and approved the manuscript.

## Code availability

The TipQuant code will be publicly released upon publication.

## Data availability

All raw video data used in the manuscript are available on Dryad through https://doi.org/10.5061/dryad.dr7sqvbd2 (Available after manuscript acceptance)

## Supporting Information

Supplemental Method S1. Contour segmentation

Supplemental Method S2. Tip detection

Supplemental Method S3. Measurement of mean fluorescence intensity and generation of PM kymograph

Supplemental Figure S1. Simultaneously analyses of ROP activity and PM Ca^2+^ influx in pollen tubes. (additional data related to Figure 5)

