## supplemental method for "TipQuant: A robust algorithm for quantitative analysis of spatiotemporally dynamic activities in tip-growing cells"

**Supplemental Method S1. Contour segmentation**

**To increase robustness of contour segmentation, we employed two pre-processing steps detailed below:**

Gamma correction

The gamma correction is very useful considering we are working with grayscale images. This operation enhances the contrast in an image, making bright pixels look brighter and dark pixels look darker. It can help a lot; from highlighting the tip-growing cell, to clear background noise. To perform this operation, we use the classical formula from https://en.wikipedia.org/wiki/Gamma_correction, which is the inverse transfer function:

$$V_{out}=V_{in}^{\frac{1}{\gamma}}$$

where V_in_ is scaled down to values between [0, 1] and V_out_ scaled up to values between [0, 255]. We noticed that a value of $\gamma=1.5$ is generally enough for most of the images.

Gaussian blur

Once the gamma correction is done, we can proceed further into the Gaussian blur (or Gaussian filter), which helps smoothing the shape of the cell and removing small noise around its border. For the implementation, we simply follow the one from OpenCV (cv.GaussianBlur()) described in the documentation:

$$G\left( x,y \right)=Ae^{\frac{-\left( x-\mu_{x} \right)^{2}}{2\sigma_{x}^{2}}+\frac{-\left( y-\mu_{y} \right)^{2}}{2\sigma_{y}^{2}}}$$

where A is the amplitude of the Gaussian, μ is the mean and σ^2^ represents the variance (per each of the variable x and y). We found that a kernel size of 7, meaning a 7x7 matrix, and a value of 5 for sigmas produces good results.

#### **Binary cell mask using a global threshold method**

Next, we apply a threshold to the image to get a binary mask of the cell. The value of the threshold is computed beforehand and for the whole video. As the overall fluorescence intensity is varying through the video due to photobleaching of fluorophores, using an adaptive threshold technique for each frame would result in having a higher threshold when the initial frame is brighter than later frames, which might result in some holes inside the imaged cell when thresholding. Thus, we first skim the whole video, using the Otsu adaptive thresholding method from OpenCV, and then keep the minimum computed threshold as the value for a global threshold method during the pre-processing.

#### **Filling holes in the binary cell mask**

Once the threshold is applied during pre-processing, a binary mask for the tip-growing cell is extracted. As in many circumstances, the internal region of the cell is less bright than the border, it is possible that the binary mask does not cover the entirety of the cell. This processing consists of filling potential holes that might persist after the threshold operation. Instead of relying on morphological operations like closing, which might result in imprecise shape of the cell, we prefer to use our own function to fill the holes. First, we determine the orientation towards which the tip-growing cell might be growing by counting the number of pixels located on the borders of the frame, e.g., if there are more pixels located on the top or bottom line of the video, it would mean that the cell is growing vertically. Once we know that the cell is either growing vertically or horizontally, we can proceed to fill the missing parts. For this, we span the frame in the same orientation as the cell, i.e., if the cell grows vertically, we span the Y-axis of the frame, and vice-versa. During the spanning, we find the minimum and maximum indices where pixels are non-negative on the other axis. Considering our example on the Y-axis, the minimum and maximum indices should be on the X-axis, and they would represent the two sides of the cell. Finally, we replace the value of all the points between those two points by 255. The method is straightforward once we know the orientation of the cell, however as of now it is not robust against cells which turn drastically, ie. cells turning more than 90° or making a U-turn.

Contour adjustment

This operation consists of computing the mean of all the pixels in the frame that are on the contour of the binary mask, which we will call "contour mean value" from now on. The goal of the operation is to improve the contour mean value through erosion of the mask until we reach a maximum, i.e., until our contour fit the tip-growing cell more precisely.

More specifically: a maximum value for the contour mean value is computed as our baseline. Then, in a loop, we erode the mask the OpenCV function cv.erode(), with a small 3x3 kernel filled with ones in order to "improve" the *contour mean value* on the eroded the mask. If no improvements are observed on the first iteration, the original mask is returned, and no adjustments are done. But if we find a higher contour mean value, by a difference of at least 5, after one erosion, we continue through the loop by varying the *iteration* parameter of cv.erode() with the iteration number of the loop. In other words, we steadily increase the number of iterations of erosions with our loop, for a default maximum of 10 erosions, until we find the "best fit" for a contour. During the loop, we keep track of the value gaps, i.e., the differences observed between the contour mean value of two successive iterations. The "best fit" is such that the gap must be at least twice greater than the previous gap. Once the gain between two iterations does not satisfy this condition, we stop the loop and set the maximum number of iterations at this value for the rest of the video. As an example, if only 4 iterations in the loop (i.e., for the erosion) are needed on the first frame, then there will not be more than 4 iterations on the next frames, though there can be less. If the 10^th^ iteration is reached without finding any improved contour mean value, that means the current contour already fits the cell well enough, so no adjustment is computed. We could also assume the video might be very dark or noisy.

### **Contour smoothing**

To add smoothing, we compute the average between two consecutive contours with the cv.distanceTransform() function from OpenCV. In some cases, where some parts of the cell might disappear from the video because it is growing vertically in the space, instead of computing the average contour we might compute the union of the consecutive contour, to keep in "memory" the shape of the cell throughout the whole video.

### **Contour parameterization and characterization**

The spline parameters are extracted with the function splprep() from the interpolate module of SciPy.

To each contour point is associated a tangent vector, a normal vector, which we define as the rotation of the tangent vector, and a curvature coefficient, computed by exploiting that the normal vector is proportional to the derivative of the tangent vector, see https://en.wikipedia.org/wiki/Curvature for standard theory of the curvature of plane curves. The vectors characterizing the contour are represented in a normalized vector space, and not in the frame coordinate system. The tangent vector is obtained with the function splev() from the interpolate module of SciPy.

$$T\left( s \right)=\left( \begin{matrix} x^{'}\left( s \right) \\ y^{'}\left( s \right) \end{matrix} \right);N\left( s \right)=\left( \begin{matrix} -y^{'}\left( s \right) \\ x^{'}\left( s \right) \end{matrix} \right);k\left( s \right)=\left| \left| T^{'}\left( s \right) \right| \right|=\sqrt{x^{''}\left( s \right)^{2}+y^{''}\left( s \right)^{2}}$$

**Supplemental Method S2. Tip detection**

#### **Growth region determination by identifying displacements of cell contour between frames**

The difference between two masks from two consecutive frames is very often too small to provide an accurate estimate of the tip, we therefore introduced an interval parameter, which we call "step", indicating how many frames in the future we must look at. In other words, to observe the growth region at frame t_2_ with a step parameter D_step_, we should take the difference of the masks from the frames at t_2_ and t_2 + Dstep_. From the growth region obtained by the difference, we can retrieve the common part from it and the contour of the frame t_2_ to determine where the growth "happens" on the contour. The value of this parameter varies according to the real growth speed of the tip-growing cell. A fast growth cell requires in a small step value, and reciprocally. It is a parameter that needs to be chosen carefully and manually, after observing and judging the growth speed of the cell in the measured video. Of course, having such a parameter, implies that no analysis is done for the last D_step_ frames of a video.

### **Determination of tip position**

Tip position is determined by a sliding window around the contour of the membrane area of the cell. On each point of the contour, the sliding window computes an average dot product (inner product) between the normal vectors inside the sliding window and the vector of the direction. For sake of completeness, we remind that if the normal and direction vectors have Euclidean coordinates $N=(N_{x},N_{y})$ and $d=(d_{x},d_{y})$, then the dot product is defined as $N_{x}d_{x}+N_{y}d_{y}$, and represents the cosine of the angle between $N$ and $d$ (as the vectors have both length equal to one). Hence, the dot product is 0 when the vectors are orthogonal, and equal to one when they are identical, and to -1 when they are opposite directions. The average dot product is then associated with the central point of the sliding window. The point with the maximal value for the average dot product is considered the tip.

As shown in Figure 1B, the window of a size determined by the user, slides along the membrane on each pixel of the contour. For each step, we compute the weighted average of the dot products between the normal vector of each pixel and the direction vector. The normal vectors are weighted by the exponential of their displacements. Before being averaged, the resulting dot products are also weighted by the distance of the corresponding point to the previous tip location.

$$\frac{1}{n}\sum_{j}^{n} dist\left( j,t \right)*\left[ \left( e^{disp\left( j \right)}*N\left( j \right) \right)\cdot d\left( j \right) \right]$$

where n is the number of points considered in the contour and dist(j, t) is the normalized distance between the tip t and the point j. The distances are obtained beforehand for all pixels considered by the sliding window (including those outside the frontier of the growth region) and then normalized between 0 and 1 as a factor. A distance of 1 means the point is the closest point to the previous tip, and 0 the farthest point to the previous tip.

The displacements vectors are scaled by the exponential function to give more importance to normal vectors with a higher collinearity to the direction vector. Whereas, the distance factor, weighs the importance of a dot product. Considering a sliding window groups multiple vectors, the more the sliding window is close the previous tip location, the higher the distance factors of all the vectors inside will be. Hence, the closer the sliding window gets from the old tip location, the more its vectors will be considered in the resulting value.

### **Tip correction**

We use the DBSCAN algorithm from Scikit-learn to spot badly placed tips and correct their position according to the next or previous known valid tips. The parameters e and minPts are subtle to tweak but we found that e = 2, which is then converted into a distance in mm, and minPts = 15 generalized well enough for most of the videos.

**Supplemental Method S3.** **Measurement of mean fluorescence intensity and generation of PM kymograph**

#### Mean fluorescence intensity in user-defined cytosolic ROIs

In cytosolic ROI A, B and C, the depth acts as the distance in µm from the tip location to the "center" of the ROI. In those regions, the center is defined as the middle point between the two sides of the cell. Finally, the radius in ROI-C is the radius of the circle in the region, measured in µm (Figure 3A).

Mean fluorescence intensity on user-defined plasma membrane (PM)

Two parameters are given in input by the user: the length and the thickness of the membrane. The length tells how far apart from the tip the membrane should spread, and the thickness describes how deep towards the inside the line should stretch along the contour (see Figure 3B). As an example, a length of 20 µm would mean than the membrane spreads of 10 µm to the left and to the right of the tip. Once the list of indices is constructed, we can construct the membrane mask with the OpenCV function cv.polylines(), which will return the corresponding closed mask of the membrane with the desired thickness given in input to the OpenCV function. Again, we use the parameter *pixel size* to approximate the length in µm of the membrane. By doing so, the measured fluorescence intensity is in the scale of the pixel values so, dark videos are expected to have a lower fluorescence intensity, as opposed to bright videos. Therefore, the quality of the video can have a strong impact on the fluorescence intensity measurement.

#### PM kymograph generation --Membrane intensity distribution

We span the normal vectors of the membrane backward to measure intensity along the normal direction for each point of the membrane contour. The length of the vectors is determined by the membrane thickness parameter given in input by the user. Intensities on the spanned vectors are then averaged for each membrane contour point. Additionally, measures are interpolated to lie on the given scale so that we can compare measures from frame to frame. We can then construct a heatmap to observe the results. The intensities values here are the values of the pixels in the original frame.
